# Activation of Src-family kinases orchestrate secretion of flaviviruses by targeting mature progeny virions to secretory autophagosomes

**DOI:** 10.1101/2020.01.12.903062

**Authors:** Ming Yuan Li, Trupti Shivaprasad Naik, Lewis Yu Lam Siu, Oreste Acuto, Eric Spooner, Peigang Wang, Xiaohan Yang, Yongping Lin, Roberto Bruzzone, Joseph Ashour, Sumana Sanyal

## Abstract

Among the various host cellular processes that are hijacked by flaviviruses, very few mechanisms have been described with regard to viral secretion. Here we investigated how flaviviruses exploit the Src family kinases (SFKs) for exit from infected cells. We isolated three members of the SFK family – Src, Fyn and Lyn – that were specifically activated during secretion of Dengue and Zika or their corresponding virus like particles (VLPs). Pharmacological inhibition or genetic depletion of the SFKs blocked virus secretion, most significantly upon Lyn-deficiency. Lyn^-/-^ cells were severely impaired in virus release, and were rescued when reconstituted with wild-type Lyn, but not a kinase- or palmitoylation-deficient Lyn mutant. We further established that Lyn, via its palmitoylation-dependent membrane association, triggered post-Golgi virus transport in specialised Rab11 and Transferrin receptor positive organelles resembling secretory autophagosomes, and distinct from conventional exocytic vesicles. In the absence of Lyn activity or its aberrant membrane association, virions were sorted into the lysosomal pathway for degradation. This mode of export was specifically triggered by processed, and mature, but not by furin-resistant virus particles, and occurred with significantly faster kinetics than the conventional secretory pathway. Our study therefore charts a previously undiscovered Lyn-dependent exit strategy, triggered by flaviviruses in secretory autophagosomes that might enable them to evade circulating antibodies and dictate tissue tropism.

## Introduction

Dengue and Zika represent two of the major mosquito-borne flaviviruses that collectively have huge health implications worldwide ^1–3^. Dengue infects approximately 400 million people annually, often causing severe pathologies such as vascular endothelial leakage. ZIKV too has emerged as a global threat with recent outbreaks linked to serious neuro-developmental complications in children and Guillain Barré syndrome in adults ^4^. Vaccines and therapeutic options for these viruses are currently unavailable, along with limited knowledge on the underlying mechanisms of pathogenesis and viral manipulation of host cell biology.

Dengue and Zika viruses exhibit significant overlap in their genome organization, intracellular life cycle and exploitation of host cellular processes ^5–7^. Both rely on specific interactions with host factors that are necessary for viral entry, replication, assembly and release ^8^. Although several genetic screens have uncovered host dependency factors for viral entry and replication ^9^, few studies have investigated mechanisms underlying assembly and release of progeny virions. Infection is accompanied by induction of lipophagy ^10, 11^, followed by massive membrane reorganization and formation of assembly sites that appear as membrane invaginations along the ER ^12^. Immature virions bud into the ER lumen, and traffic along the cellular secretory pathway from the ER through the Golgi complex, where they undergo maturation to form infectious particles prior to exiting the cell via as-yet undefined routes.

To address mechanisms of virus release, we previously established a cell-line that stably expresses the Dengue structural proteins pre-membrane (prM) and envelope (E), and constitutively secretes non-infectious recombinant virus like particles (VLPs) ^13, 14^. Transport of these VLPs mimic the intracellular trafficking characteristics of infectious Dengue particles – thus providing a convenient method to identify the molecular determinants of Dengue secretion without confounding effects from viral entry and replication. Using this tool, we determined that interaction between Dengue prM and cellular KDEL receptors (KDELRs) facilitated ER-to-Golgi transport of newly formed Dengue particles ^13^.

KDELRs consist of seven transmembrane segments and have a similar topology as that of G-protein coupled receptors (GPCRs). Analogous to GPCR-signaling pathways, KDELRs are also able to recruit heterotrimeric G-proteins to activate Src-family kinases (SFKs) – in particular, Src at the Golgi complex. The KDELR-Src signaling axis is crucial for sensing increased protein flux from the ER to the Golgi apparatus, and activating anterograde transport to maintain inter-organelle homeostasis^15,16^.

Since Dengue exploits KDELRs to arrive at the Golgi, we hypothesized that SFK-dependent signaling is very likely triggered during anterograde transport of progeny virions. We therefore screened for activated SFKs in Dengue-infected cells. The SFK family comprises nine members that are ubiquitously expressed in different combinations in all tissues ^17^. Of these, we identified three – Lyn, Fyn and Src that were heavily phosphorylated in their activation loop (Y420 for Fyn, Y397 for Lyn and Y419 for Src) in infected and VLP-secreting cells. Inhibiting their activation by pharmacological means or individual gene-depletions blocked virus secretion. Among the three SFKs, Lyn-depletion displayed the most significant defect. We therefore created Lyn^-/-^ cells using CRISPR/Cas9 and reconstituted with its wild-type and mutant variants. Lyn^-/-^ cells were severely impaired in secretion of progeny virions and VLPs. Transport from the ER to the Golgi remained unaffected, but failed post-Golgi, and were sorted for lysosomal degradation instead. This defect could be rescued by reconstituting a wild-type variant of Lyn in the Lyn^-/-^ background. On the other hand Lyn mutants that were deficient in: (a) kinase activity or (b) N-terminal palmitoylation, failed to rescue this defect. Using biochemical and imaging analyses, we established that Lyn, via its palmitoylation-dependent membrane association, triggered post-Golgi virus transport in specialised organelles that were Rab11 and Transferrin receptor positive, indicative of secretory autophagosomes. This mode of Lyn-dependent transport was specifically triggered by processed, mature virions, but not by protease-resistant immature ones that were instead exported with significantly slower kinetics, and in a Lyn-independent manner. Interestingly, expression of Lck, which belongs to the same SFK subfamily as Lyn, was able to rescue the secretion defect in Dengue-infected Lyn^-/-^ cells, whereas acylation-defective Lck mutants failed to do so, reminiscent of Lyn activity. The mechanism of flavivirus secretion has long been speculated; our study uncovers a novel Lyn-dependent exit strategy triggered by flaviviruses that might enable them to evade circulating antibodies and dictate tissue tropism.

## Results

### Flavivirus infection activates Src family kinases

A pulse of protein flux from the ER to Golgi compartments is known to activate a KDELR-Src-dependent signalling cascade that is necessary to maintain inter-organellar trafficking. Since Dengue progeny virions bind to KDELRs to arrive at the Golgi, we hypothesised that an analogous SFK-dependent signalling is likely triggered during infection to enable virus transport. To test this hypothesis, we measured activation of SFKs in mock- and Dengue virus-infected cells using antibodies that specifically recognise the phosphorylation at their activation loop. SFKs displayed >5-fold increase in their phosphorylation status in infected cells measured at varying MOI (0.1, 1, 5 or 10) in two different cell lines (BHK21 and Vero) (**Figure 1A**). We verified this phenomenon in Zika virus infected cells, which is a closely related flavivirus. Phosphorylation of SFKs increased both in a time and dose-dependent manner (**Figure 1B**).

**Figure 1.**
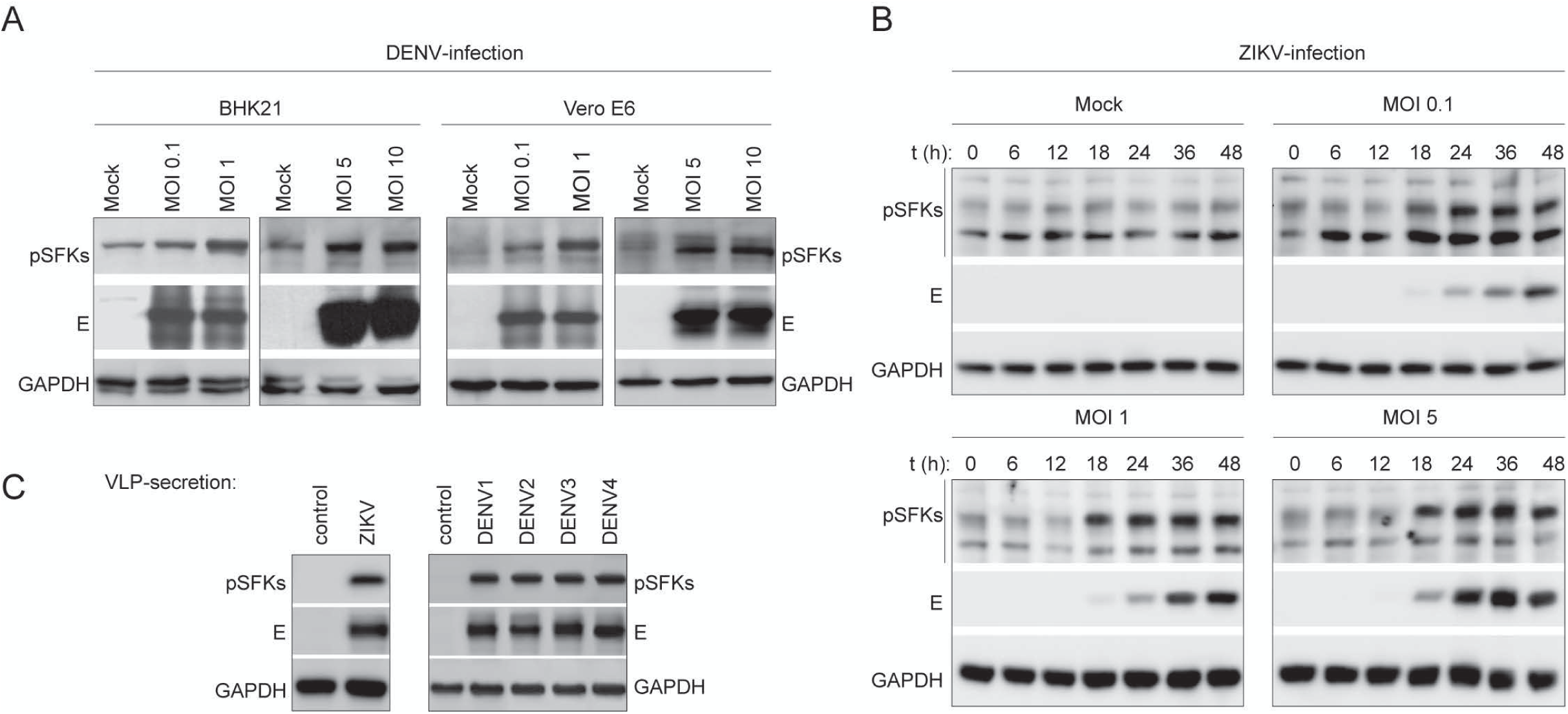
Flavivirus infection triggers activation of Src-family kinases. (A) Two different cell types (BHK21 and Vero E6) were infected with Dengue virus at varying MOI (0.1, 1.0, 5 and 10). Lysates prepared from infected cells were immunoblotted with antibodies against phosphorylated SFKs and anti-flavivirus envelope 4G2 antibodies to detect virus particles. Gapdh was used as loading control. **(B)** Cells were infected with Zika virus at varying MOI of 0.1, 1 and 5. At different time intervals post infection lysates prepared from mock and infected cells were immunblotted with anti-phospho SFK antibodies to measure activation and 4G2 antibodies to detect virions. **(C)** Cells constitutively secreting either Zika (*left panel*) or Dengue (*right panel*) VLPs were immunoblotted to detect phosphorylated SFKs. All images are representative of atleast 3 independent experiments.

To focus exclusively on virus release, we took advantage of a VLP-secretion model system. These cells stably express the viral E and prM proteins (from Dengue serotypes 1-4, or Zika), and constitutively secrete VLPs, thus mimicking virus budding, maturation and release ^14^. Here too, phosphorylation of SFKs was significantly enhanced (>5 fold) in cells secreting VLPs for all Dengue serotypes as well as Zika, as compared to their parental control cells (**Figure 1C**). These results support our hypothesis that intracellular transport of progeny virions or VLPs, irrespective of their serotype, triggers activation of cellular SFKs.

### Identification of SFK members involved in flavivirus infection

There are at least nine members of SFKs expressed in different combinations in all mammalian cells ^17^. To identify those that are activated upon Dengue infection we immunoprecipitated phosphorylated SFKs using anti-pSFK antibodies from cell lysates prepared from mock or Dengue virus-infected BHK21/Vero E6 cells. Phosphorylated SFKs and associated cellular factors were isolated, resolved by SDS-PAGE, and detected by silver staining (**Figure 2A**). The lanes were sliced into 2mm sections and subjected to trypsin digest for identification by mass spectrometry. We identified three members – Src, Fyn and Lyn, and several SFK-regulated substrates in Dengue-infected samples (**Figure 2B**). Among the co-immunoprecipitating proteins several were components of the secretory pathway, ER/Golgi resident proteins and vesicular transport machinery including the KDELRs, which we had previously characterized as important for dengue secretion ^13, 14^.

**Figure 2.**
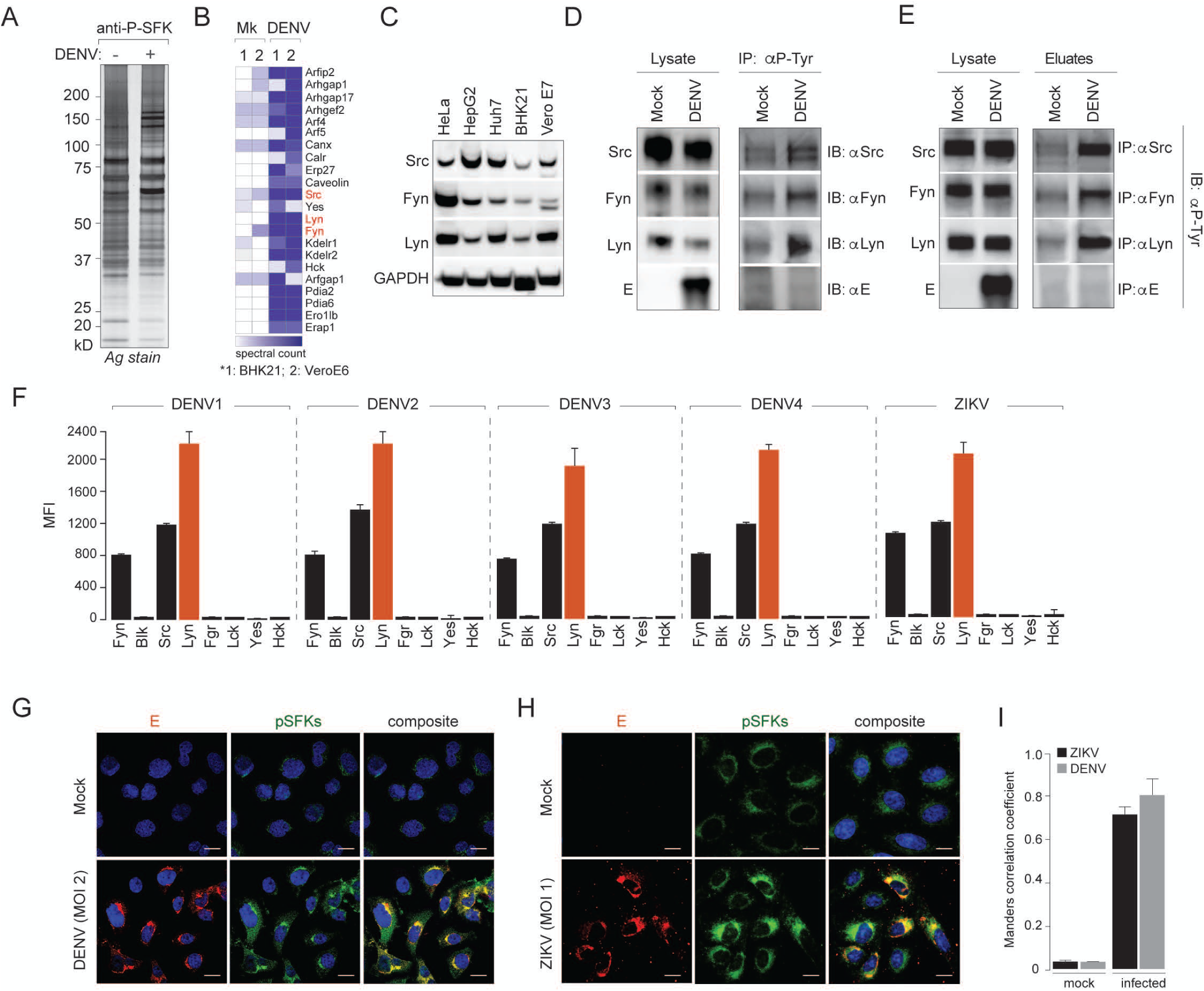
Flavivirus infection triggers activation of three specific SFKs. (A) Large scale immunoprecipitation of activated SFKs was performed on anti-phospho-SFK antibodies from mock and Dengue-infected BHK21 and Vero E6 cells. Isolated proteins were resolved by SDS-PAGE and detected by silver staining. **(B)** Entire lanes on gels were sliced into 2mm sections and subjected to trypsin digest. The peptide mix generated was processed and analysed by an LTQ Orbitrap mass spectrometer. **(C)** Protein expressions of Lyn, Fyn and Src kinases identified by mass spectrometry were validated in Dengue susceptible cell types. **(D)** Activation of Lyn, Fyn and Src upon Dengue infection was measured by immunoprecipitating first on anti-phospho-Tyrosine antibodies and immunoblotting with specific antibodies and **(E)** a reciprocal immunoprecipitation on specific anti-SFK antibodies followed by immunoblotting with anti-phospho-Tyrosine antibodies. **(F)** Activation of SFKs was measured in lysates prepared from Dengue-infected cells with the Milliplex MAP 8-plex assay kit using the Luminex system as read-out, following the manufacturer’s protocol. **(G)** Colocalisation of Dengue and SFKs was visualised by 4G2 antibodies and anti-phospho-SFKs. **(H)** Same as (G) in Zika-infected cells. **(I)** The extent of Dengue and Zika colocalisation with activated SFKs was measured by Manders correlation coefficients.

The three SFKs displayed high levels of expression in Dengue susceptible cell lines (**Figure 2C**). To confirm their activation, we immunoprecipitated phosphorylated proteins from mock and Dengue virus-infected cells using anti-p-Tyrosine-antibodies. Eluates from immunoprecipitated material were analysed by Western blotting with specific antibodies against the selected kinases - Src, Fyn and Lyn (**Figure 2D**). Although expression levels of total SFKs were comparable in mock and infected samples, a significant increase was noted in their phosphorylated form in eluates from Dengue virus-infected samples compared to mock (**Figure 2D**). We also employed a reciprocal strategy where kinases from mock and Dengue-infected samples were first immunoprecipitated on specific antibodies followed by immunoblotting with anti-phospho-tyrosine antibodies, which confirmed these results (**Figure 2E**). The Dengue structural E-protein however, did not co-purify with phosphorylated SFKs, suggesting that they do not directly interact with each other. To further quantitate increases in specific SFK activation, we took advantage of the Milliplex Map 8-plex SFK activation kit using the Luminex technology, which confirmed specific activation of Lyn, Src and Fyn in lysates prepared from Dengue and Zika-infected cells (**Figure 2F**).

To visualise the subcellular distribution of pSFKs upon virus infection, we performed confocal imaging in both Dengue (**Figure 2G**) and Zika infected (**Figure 2H**) cells. Whereas in mock-infected cells, SFKs appeared to be confined to the Golgi compartments, upon virus infection, the distribution spread throughout the secretory pathway and appeared along the ER, Golgi and the plasma membranes, with significant overlap with the viral E-protein distribution (**Figure 2I**). Collectively, these data suggest that upon infection, progeny virus particles colocalise with SFKs along the secretory pathway. However, they either do not physically associate or participate in transient interactions typical of enzyme-substrate complexes.

### SFKs facilitate flavivirus production

To study the role of selected SFKs in flavivirus infection, particularly in viral transport along the secretory pathway, we investigated the effect of inhibiting SFK activity using two approaches. First, we selected a commercially available SFK inhibitor – SU6656 ^18^, which specifically blocks their activation as measured by the absence of their phosphorylated forms (**Figure 3A**). We performed an MTT assay to establish that at concentrations of SU6656 <10µM, cell viability was not affected (**Figure S1**). We selected this concentration range to test virus production and secretion of VLPs upon SU6656 treatment. Our results indicated that 5µM SU6656 was sufficient to cause ∼10-fold reduction in viral titers (**Figure 3B**) and >50% reduction in secreted VLPs resolved by gel electrophoresis (**Figure 3C**). Densitometric analyses indicated that all DENV and ZIKV VLPs displayed attenuated secretion upon treatment with 5 µM SU6656 (**Figure 3D**).

**Figure 3.**
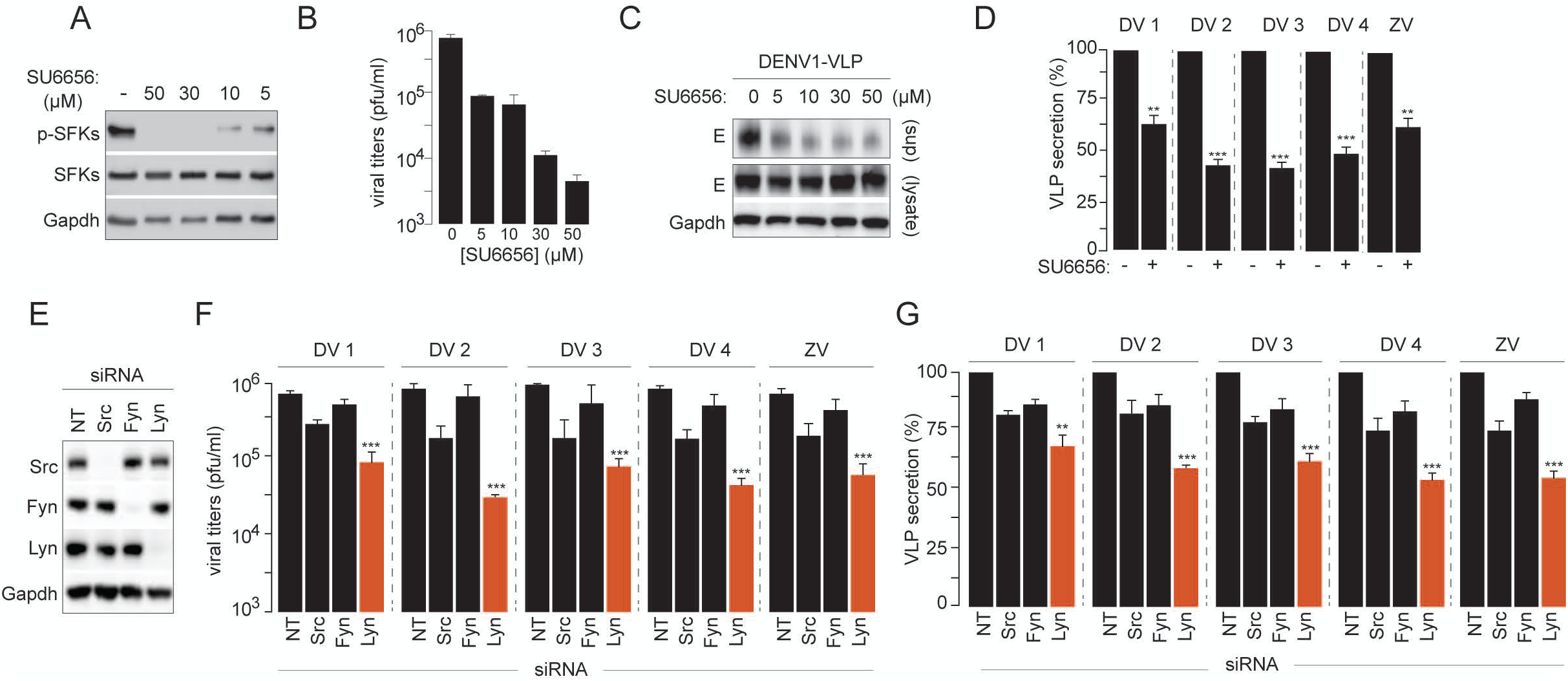
Inhibition of SFK-activation attenuates virus production as a result of impaired secretion. (A) Cells were treated with different concentrations of a selective SFK inhibitor (SU6656); phosphorylation of the activation loop was measured by immunblotting with anti-phospho-SFK antibodies. Total SFKs were detected using anti-Src rabbit polyclonal antibodies cross-reactive against all SFK members. Gapdh was used as loading control. **(B)** Cells treated with SU6656 were challenged with Dengue virus (DENV 2) at MOI 2. 36 hours post infection, supernatants from infected cells were measured for production of virus particles using plaque assay. Error bars represent mean±s.d from three independent biological replicates. **(C)** Secretion of Dengue VLPs from cells treated with varying concentrations of SU6656 was measured by immunoblotting with anti-Dengue 4G2 antibodies, in supernatants and cell lysates. Gapdh was used as loading control. **(D)** Quantitation of VLP secretion described in (C) by densitometric analyses for all serotypes of Dengue and Zika virus. Percentage VLP secreted was calculated as fraction of total intracellular VLPs normalised to untreated cells set at 100%. Error bars represent mean±s.d from three independent biological replicates; Statistical significance was calculated using Student’s T test; *** represents p<0.001. **(E)** Depletions of Lyn, Fyn and Src were performed by siRNA targeting individual kinases, and verified by immunoblotting with kinase specific antibodies. **(F)** SFK-depleted cells were challenged with different serotypes of Dengue virus and Zika. Viral titers were measured in supernatants collected from infected cells, using plaque assays. Error bars represent mean±s.d from three independent biological replicates. **(G)** Depletion of SFKs was performed in the corresponding VLP-producing cells to measure secretion, using 4G2 antibodies to detect viral structural proteins as readout. Error bars represent mean±s.d from atleast three independent biological replicates.

To genetically validate the role of SFKs in viral secretion, we first transfected VLP-secreting cells (Dengue, all serotypes and Zika) with siRNA targeting the individual kinases. Immunoblots using selective antibodies indicated significant reduction in protein expression of targeted SFKs, without altering that of the others (**Figure 3E**). Among the three kinases, depletion of Lyn had the most significant effect, with ≥10-fold reduction in viral titers for all strains. Depletion of Src had a modest effect, whereas that of Fyn did not affect virus production (**Figure 3F**). These data were reflected in the VLP-secretion assay, where a similar reduction was noted in Lyn-depleted cells (**Figure 3G**).

To investigate whether SFKs operated in a concerted manner to enable virus secretion, we tested various combinations of siRNAs against the selected kinases. Similar to individual siRNA depletions, combinations of siRNAs could also significantly reduce the expression of the corresponding kinases (**Figure S2A**). Compared to depletion of individual SFKs, combination of Lyn and Src was able to further suppress Dengue and Zika VLP secretion. However, siRNA combinations without Lyn siRNA had no significant effect on VLP secretion (**Figure S2B**). Collectively, these results suggest that Lyn plays a critical role in regulating secretion of progeny virus particles – a function reminiscent of Src-dependent protein transport from the ER to the Golgi. Virus particles are likely sorted into specific vesicular trafficking pathways that require activation and signalling by SFKs to enable their vectorial transport through the host secretory system.

### Virus secretion requires kinase activity and palmitoylation of Lyn

To confirm the role of Lyn in flavivirus infections, we generated Lyn-deleted cells using the CRISPR/Cas9 technology. Two guide RNA (sgRNA) (5’-TGAAAGACAAGTCGTCCGGG-3’ and 5’-GTAGCCTTGTACCCCTATGA-3’) targeting human Lyn mRNA were designed and cloned into the chimeric CRISPR/Cas9 vector PX459. Transfected cells were selected on puromycin and immunoblotted with anti-Lyn to verify deletion (**Figure 4A**). Lyn^-/-^ cells were also reconstituted with wild-type (rLyn) and a mutant variant of Lyn (C468A) through lentiviral transduction. Cysteine 468 is located at the C-terminus of the kinase domain, and the Lyn mutant carrying an alanine substitution at C468 impairs phosphorylation of Lyn in its activation loop, thereby blocking its kinase activity ^19^. An additional mutant with a cysteine to serine substitution was generated to prevent palmitoylation of Lyn. All of the generated cells (WT, Lyn^-/-^, rLyn, C468A, C3S) were then challenged with either the 4 serotypes of Dengue or Zika virus. We collected supernatants from mock and infected cells to measure viral titers by plaque assay. Virus production was significantly attenuated in all Lyn^-/-^ cells, and could be rescued in the rLyn expressing cells, but not in those expressing the kinase inactive C468A mutant or the palmitoylation deficient C3S mutant (**Figure 4B**). To determine whether Lyn-deficiency caused defective viral entry or replication, we measured viral RNA using RT qPCR upon Dengue and Zika infection from cell lysates 24 hours post infection, and did not detect any difference (**Figure 4C**). We also measured E-protein synthesis in metabolically labelled cells to rule out any defect in viral protein translation. Again, the abundance of newly synthesised viral structural proteins were comparable in wild-type and Lyn-deficient cells or those expressing its mutant variants, indicating that viral protein translation was not affected in the absence of Lyn activity (**Figure 4D**).

**Figure 4.**
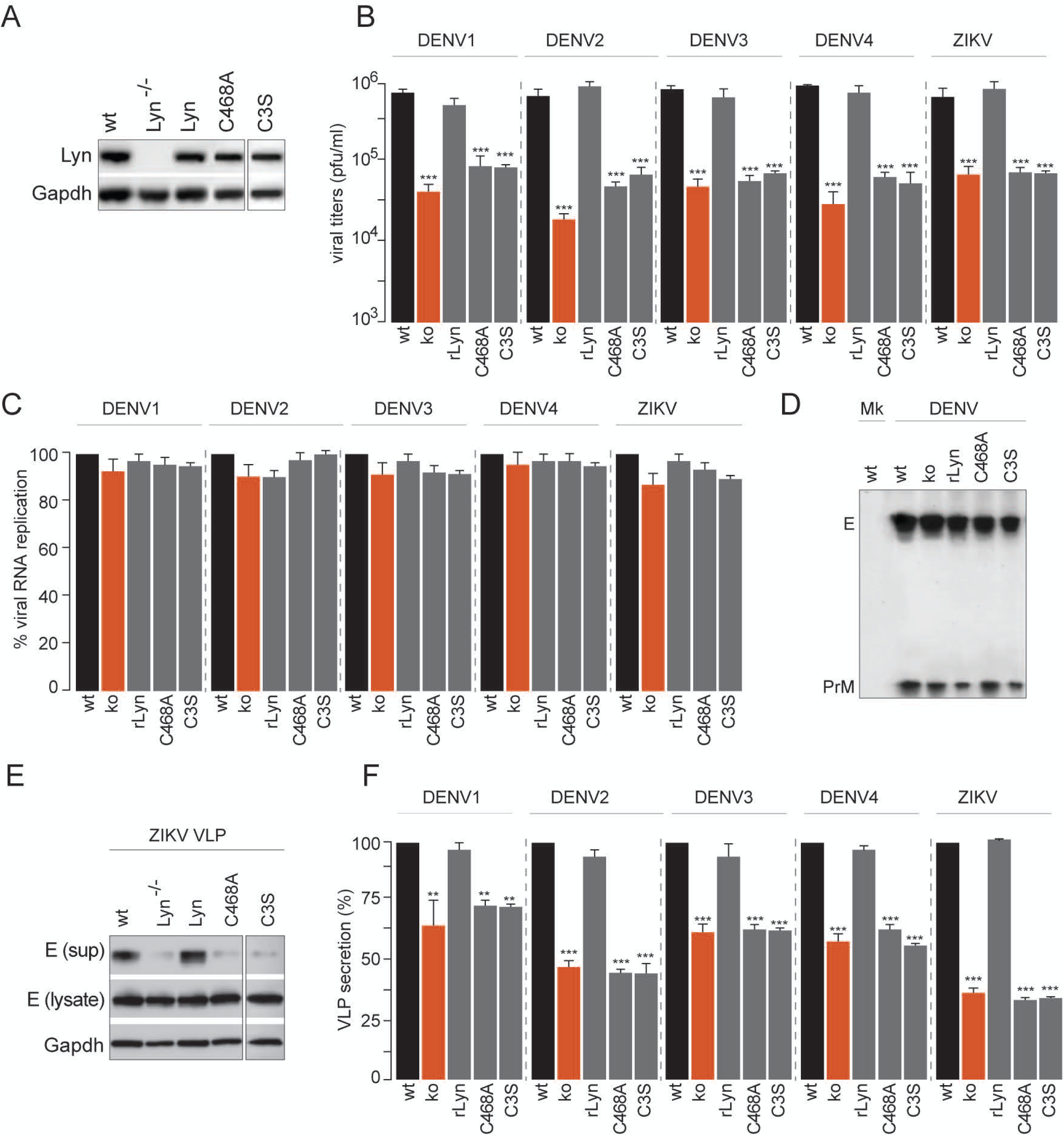
Characterisation of Lyn activity in flavivirus secretion. (A) Lyn-deficient cells were generated by CRISPR/Cas9 mediated gene deletion. Lyn^-/-^ cells were reconstituted with either (i) wild-type Lyn, (ii) C468A kinase-inactive mutant or C3S palmitoylation deficient mutant. Validation of gene deletion and expression of Lyn mutants were performed by immunoblotting in cells expanded from single clones. **(B)** Cells described in (A) were challenged with different serotypes of Dengue or Zika virus at MOI 2. Supernatants from infected cells were collected 48 hours post infection and measured for viral titers using plaque assays. **(C)** Virus replication was measured in cells described in (B) using RT qPCR. Error bars represent mean±s.d from three independent biological replicates; statistical significance was calculated using Student’s T test; *** represents p<0.001. **(D)** Viral protein synthesis in wild-type versus Lyn^-/-^ cells or those expressing Lyn mutants was determined by metabolic labeling with [^35^S]cysteine/methionine. Viral structural proteins were isolated from cell lysates at early time points on anti-E 4G2 antibodies and detected by autoradiography. **(E)** Supernatants were collected from VLP-secreting cells which were either wild-type or Lyn^-/-^, or reconstituted with (i) wild-type Lyn, (ii) C468A kinase-inactive mutant or (iii) C3S palmitoylation-deficient mutant. Concentrated VLPs were resolved by SDS-PAGE and immunoblotted with 4G2 antibodies. **(F)** VLP-secretion was quantitated for Dengue serotypes and Zika virus as fraction of total normalised to wild-type cells set at 100%.

To verify that attenuated virus production was specifically due to defective transport, we measured appearance of VLPs in the supernatants. For both Dengue and Zika, we observed a significant loss in VLP-secretion from Lyn^-/-^ cells, which could be rescued in cells expressing wild-type Lyn (rLyn) but not C468A or C3S mutants, suggesting that both its kinase activity and palmitoylation are necessary for intracellular virus transport and release (**Figure 4E, F**).

### Virus transport is blocked post Golgi in Lyn-deficient cells

To determine the underlying mechanism of Lyn-dependent virus transport, we performed biochemical and imaging analyses of virions in infected cells. First we performed pulse-chase assays to measure virion trafficking characteristics using E-protein as read-out. Mock and infected cells were radiolabelled with [^35^S]cysteine/methionine for 1 hour, and chased in cold medium. At different time intervals post infection (0, 6 and 12 hours), intracellular virions were isolated on 4G2 antibodies, resolved by gel electrophoresis and detected by autoradiography as shown in schematic (**Figure 5A**). To assess and quantitate its intracellular distribution, we treated immunoprecipitated E with either endoglycosidase H (EndoH) or PNGaseF to measure its glycosylation status as read-outs for localisation within the secretory pathway. The viral E-protein undergoes high mannose glycosylation in the ER, and remains EndoH sensitive; upon arrival at the Golgi it acquires complex glycans and becomes EndoH-resistant. In addition, the host protease furin cleaves prM to generate soluble pr and membrane bound M, which can be resolved by gel electrophoresis. The post-ER pool of E-protein was therefore calculated as the EndoH resistant fraction of total intracellular E-protein (**Figure 5B**). In the Lyn-deficient cells or those expressing its mutant variants, kinetics of ER to Golgi transport of E-protein remained unaffected, as measured by acquisition of EndoH resistance (**Figure 5C**). Secretion of virus particles is also accompanied by appearance of membrane-bound E-protein at the cell surface. To monitor the fraction of total E that reaches the cell surface we first biotinylated the surface proteins from mock and infected cells, isolated them on streptavidin beads and detected by immunoblotting with anti-E antibodies (**Figure 5D**). Contrary to ER-Golgi transport, arrival of E-protein at the cell surface was severely impaired in Lyn^-/-^ cells, or those expressing Lyn mutants (**Figure 5E**). These data indicate that Lyn regulates virus transport from the Golgi en route to the cell surface.

**Figure 5.**
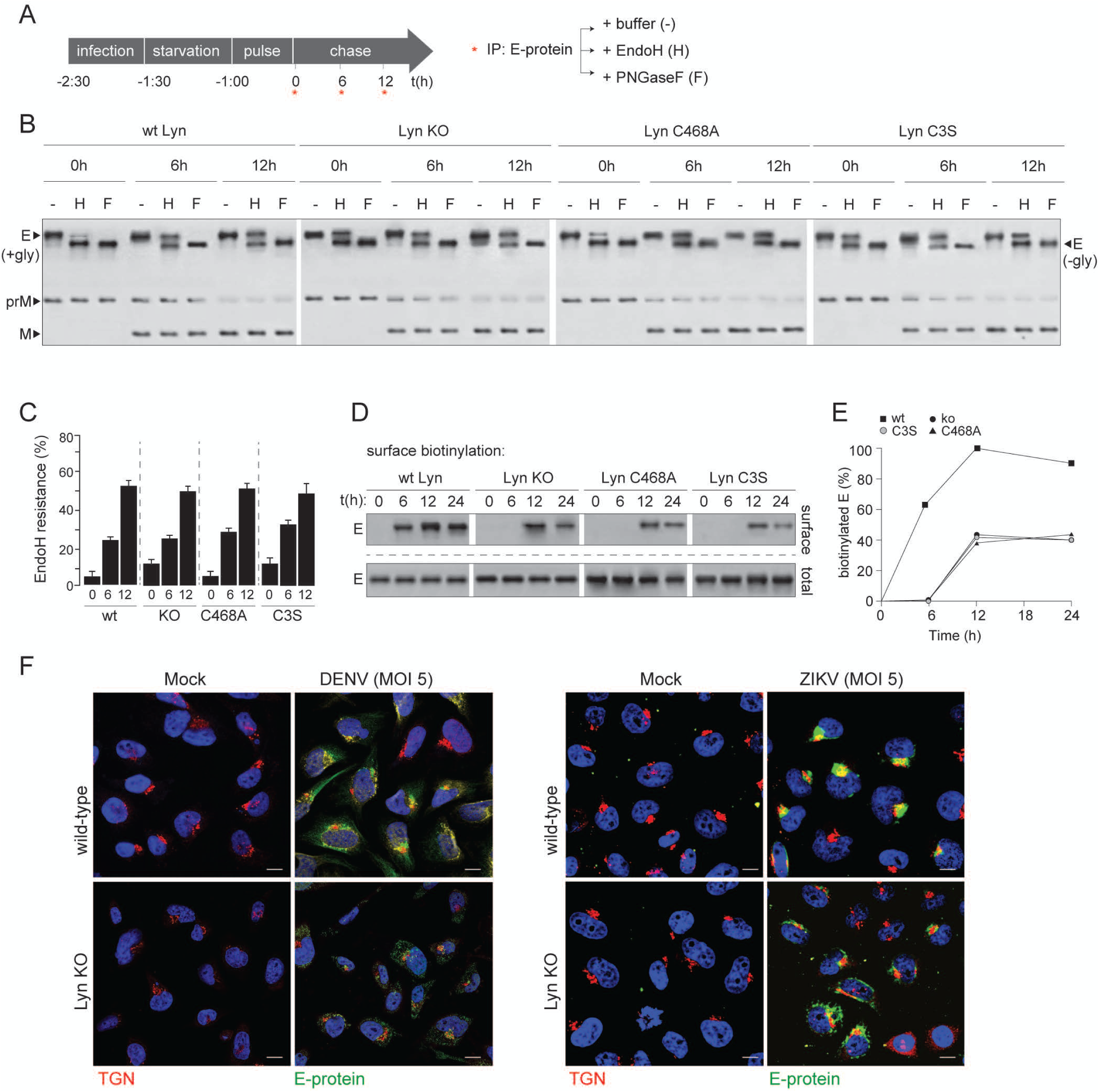
Lyn-deficiency blocks virus secretion post ER to Golgi transport. (A) Schematic of pulse-chase analyses to characterise virus trafficking. **(B)** Wild type and Lyn^-/-^ cells, or those reconstituted with either C468A or C3S mutants were infected with Dengue at MOI 5. Cells were then pulsed with [^3^5S]cysteine/methione, and chased in cold medium for 6 and 12 hours. At each time interval virions were isolated by freeze-thaw homogenisation followed by immunoprecipitation on 4G2 antibodies. Immunoprecipitated material was either treated with (i) buffer only (-), (ii) Endoglycosidase H or (iii) PNGase F. All samples were resolved by SDS-PAGE and detected by autoradiography. **(C)** Densitometric analyses were performed on autoradiograms to measure fraction of EndoH resistance as % of total E. **(D)** Surface biotinylation on intact cells described in (B) was performed by Sulfo-NHS-biotin using manufacturer’s protocol. Lysates prepared from all cells were divided into two for: (i) immunoprecipitation on streptavidin beads and (ii) total intracellular E expression. Biotin-modified E and total intracellular E from all samples were resolved by SDS-PAGE and detected by immunoblotting with 4G2 antibodies. **(E)** Quantitation of biotinylated E was performed by densitometric analyses on immunoblots from (D). Biotinylated E was calculated as % of biotinylated E in wild-type cells at 12 h set at 100%. **(F)** Mock and virus-infected cells were visualised by confocal imaging to detect intracellular localisation of viral particles using 4G2 antibodies. TGN marker was stained as control. (*Left panel*) Dengue virus infected cells; (*right panel*) Zika virus infected cells.

To visualise the block in virus secretion, we imaged distribution of viral E-protein in wild-type and Lyn^-/-^ cells, either infected with Dengue or Zika (**Figure 5F**). In wild-type cells, E-protein displayed its characteristic accumulation in membrane sites proximal to the trans Golgi network. These are believed to be modified ER membranes where progeny virions accumulate, suggestive of ER to Golgi transport as the rate-limiting step in virus trafficking ^20–22^. On the other hand, in Lyn-deficient cells, we detected aberrant E-protein distribution, which appeared to be in internal vesicles distinct from that of the trans Golgi or the modified ER membranes. Collectively our data indicate that in the absence of Lyn or in the presence of kinase-inactive or palmitoylation-deficient Lyn, synthesis, assembly and ER to Golgi transport of progeny virions remain unaffected; however, they are very likely mis-sorted into vesicles that fail to be secreted from the Golgi compartments.

### Palmitoylation-dependent membrane association of Lyn with secretory organelles determines virus secretion from post Golgi compartments

SFKs are equipped with glycine and cysteine at their N-terminus, which are typically acylated with myristoyl and palmitoyl chains respectively. S-palmitoylation has been shown to dictate the spatial distribution and trafficking characteristics of several proteins including SFKs ^23–25^. Given the detrimental effect on virus transport with palmitoylation-deficient Lyn mutant C3S (**Figure 5**), we aimed to determine how acylation affected virus secretion. We radiolabelled wild-type cells with [^3^H]myristoyl or [^3^H]palmitoyl CoA, and either mock-treated or infected cells with Dengue virus. (**Figure 6A**). Individual kinases were isolated on selective antibodies and their acylation detected by autoradiography. Fyn acquired myristoyl and palmitoyl chains in both mock and infected samples; Src did not undergo any acylation regardless of infection; whereas Lyn displayed palmitoylation in virus-infected cells.

**Figure 6.**
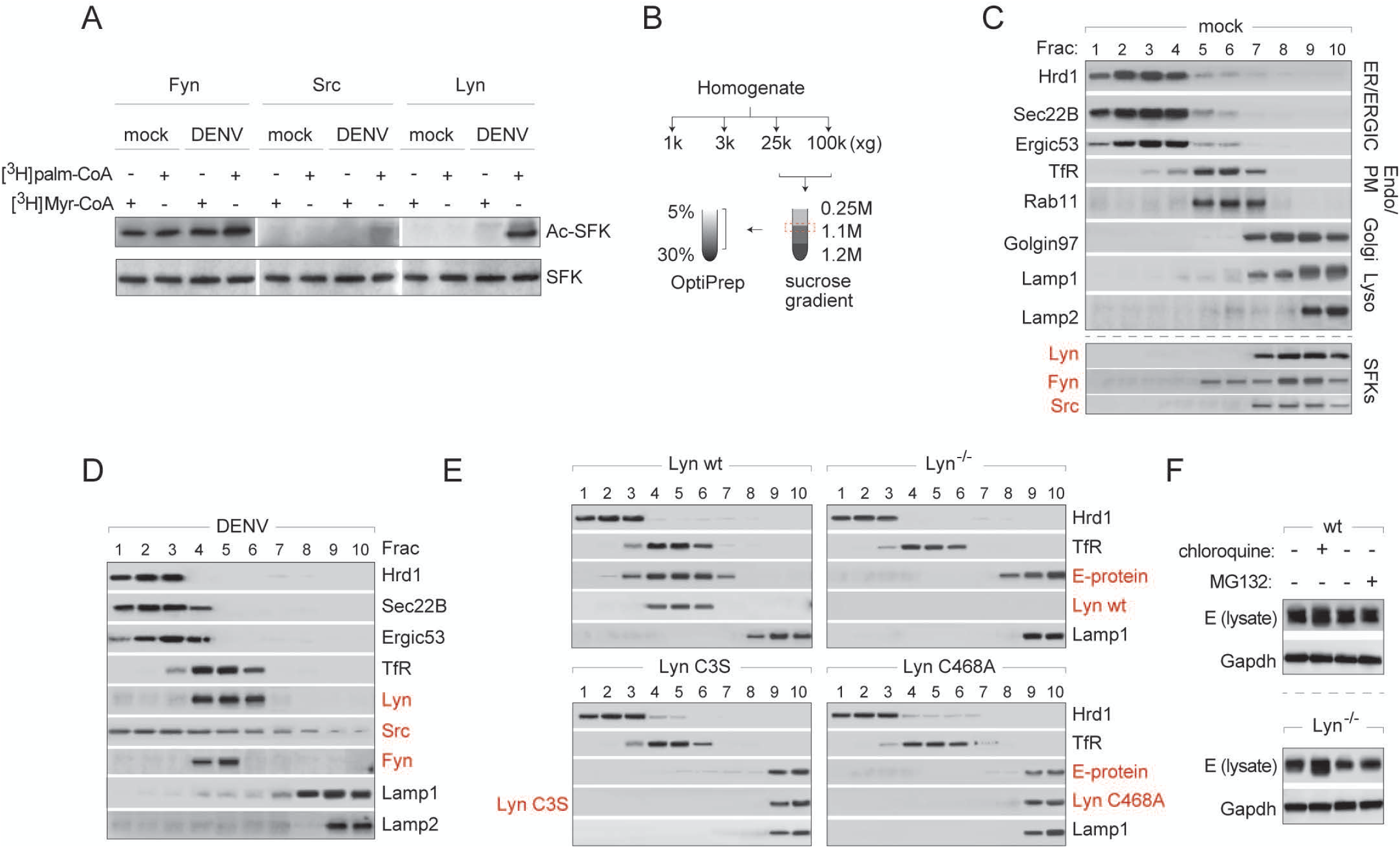
Lyn-dependent virus transport requires its palmitoylation and kinase activity. (A) Wild-type cells were radiolabelled with [^3^H]myristoyl or [^3^H]palmitoyl CoA and either mock or virus-infected. Lyn, Fyn and Src were immunoprecipitated, resolved by SDS-PAGE and detected by autoradiography. Total expression levels were measured by immunoblotting in cell lysates. **(B)** Schematic for biochemical fractionation to isolate intracellular compartments of the secretory/endolysosomal compartments. **(C)** Separation of organelle markers to verify membrane fractionation using schematic in (B). **(D)** Intracellular distribution of organelles and the three SFKs was measured in wild-type cells upon Dengue virus infection using the biochemical separation strategy described in (B). **(E)** Intracellular distribution of progeny virions (E-protein) and Lyn variants were measured in wild type and Lyn^-/-^ cells or those expressing C468A or C3S Lyn mutants, using the biochemical separation strategy described in (B). **(F)** Wild-type and Lyn^-/-^ cells were dengue-infected and either (i) untreated, (ii) chloroquine treated or (iii) MG132 treated. Expression of viral E protein was measured by immunoblotting with 4G2 antibodies.

To determine their subcellular distribution, we performed biochemical fractionations of organelles following the schematic as shown (**Figure 6B**). Modified from a previous study ^26^ this strategy can effectively enrich for membranes of the secretory and endolysosomal compartments (**Figure 6C**). In mock-infected cells, Src and Lyn co-migrated primarily with Golgi markers, whereas Fyn appeared in both the Golgi and PM fractions, in line with previous reports ^24^. However, upon infection in wild-type cells, Src displayed a broad distribution over all membranes with modest enrichment in the ERGIC compartments. On the other hand, Lyn and Fyn migrated with the Transferrin receptor and Rab11, both representative of recycling endosomes/secretory organelles depending on cellular physiology ^27^ (**Figure 6D**).

We also measured subcellular distribution of mutant Lyn variants and viral E-protein using this fractionation strategy to determine whether compartmentalisation was perturbed in Lyn-deficient cells. Both Lyn C468A and C3S mutants displayed aberrant distribution, with enrichment in lysosomal compartments in Dengue-infected cells (**Figure 6E**). Distribution of E was similarly affected. Instead of co-sedimenting with Rab11 and TfR markers as observed in wild-type cells, in Lyn^-/-^ cells or those expressing Lyn mutants, E-protein was enriched in the lysosomal compartments (**Figure 6E**).

We hypothesised that loss of Lyn palmitoylation or kinase activity resulted in defective vesicular trafficking and inhibited virus secretion from the Golgi. Accumulation of viral particles in the Golgi would therefore result in mis-sorting into lysosomes for degradation. To test this hypothesis, we blocked lysosomal degradation in wild-type and Lyn^-/-^ cells by treating with chloroquine. MG132-treated cells were processed in parallel to block the proteasomal degradation pathway. We recovered significantly more viral E-protein from Lyn^-/-^ cells compared to wild-type upon blocking lysosomal degradation (**Figure 6F**). Our data therefore suggest that activation of Lyn and its appropriate membrane targeting through palmitoylation maintains anterograde transport of progeny virus particles. Treating cells with 2-bromopalmitate (2-BP), a selective inhibitor of palmitoylation, resulted in significant loss of VLP secretion, corroborating these results (**Figure S3A, B**).

### Biogenesis of virus-triggered secretory autophagosomes is dependent on Lyn and proteolytic-processing of virions

To determine how progeny virions/VLPs triggered Lyn-dependent export via secretory organelles, we determined whether the viral envelope proteins could regulate the biogenesis of these secretory organelles. To do so, we generated stable cells secreting recombinant VLPs carrying a furin-resistant prM (H98A) and another carrying a second mutation (E E62K) that renders it furin-sensitive as reported previously ^28^ (**Figure 7A**). We purified [^35^S]-labelled VLPs from supernatants and verified that these mutations conferred furin-resistance and sensitivity respectively as detected by autoradiography.

**Figure 7.**
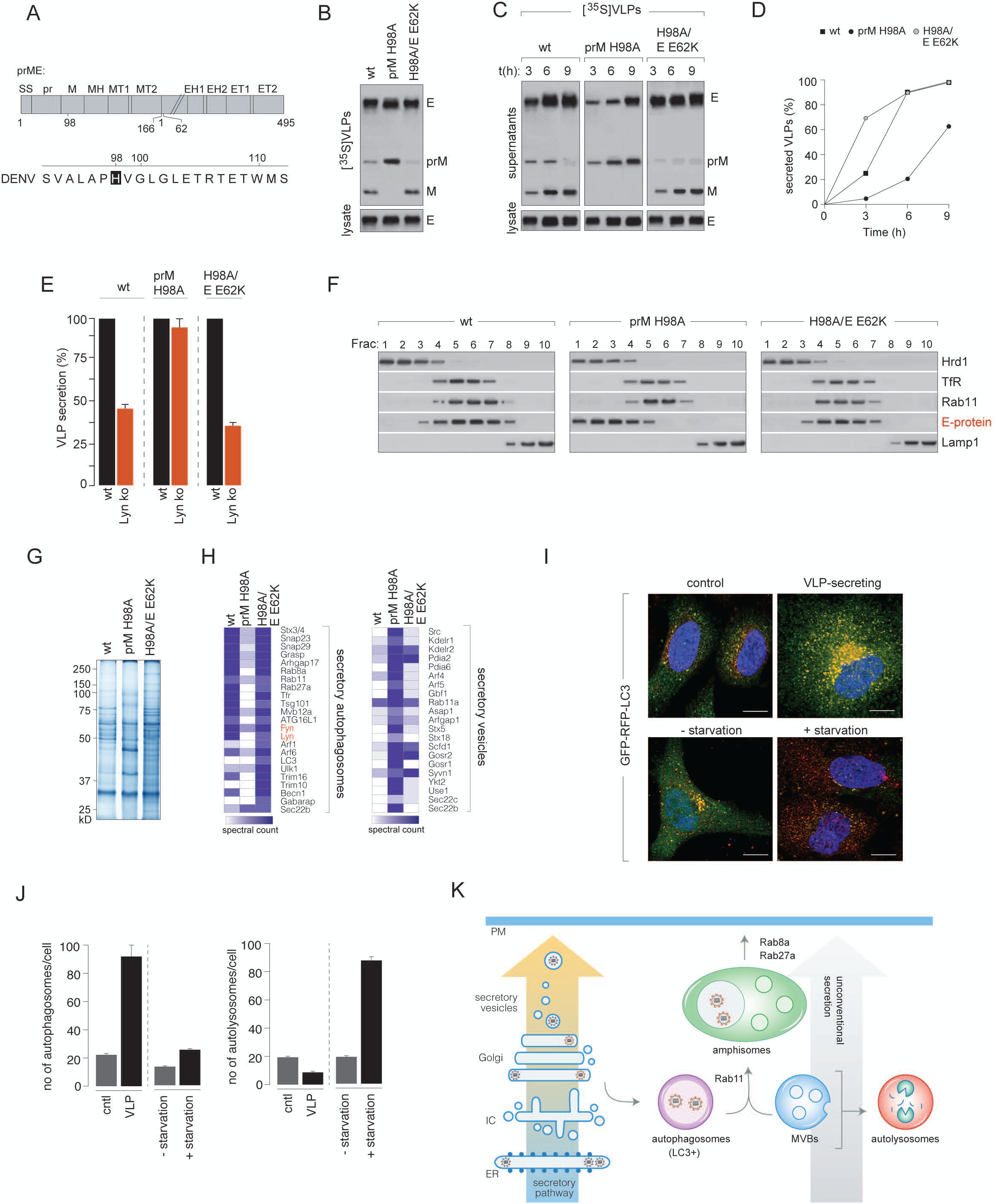
Lyn-dependent virus transport from the Golgi is specifically triggered by processed, mature virions. (A) Schematic of prME showing H98A that renders it furin-resistant and a second revertent (E E62K) that rescues furin sensitivity. **(B)** Radiolabelled VLPs, either wild-type or carrying a prM H98A mutation or a double mutation of prM H98A/E E62K were purified from supernatants of [^35^S]cysteine/methionine labeled cells. VLPs were resolved by SDS-PAGE and detected by autoradiography. **(C)** Time courses of secretion for wild-type, H98A and H98A/E62K VLP variants were measured from wild-type cells, by pulse-labelling with [^35^S]cysteine/methionine and chasing in cold medium for indicated time intervals. At each time point, VLPs were concentrated from the supernatants, resolved by SDS-PAGE, and detected by autoradiography. **(D)** Quantitation of VLP secretion was performed by densitometric analyses on autoradiogram from (C), and normalised to the wild-type sample set at 100%. **(E)** VLP secretion for wild-type, H98A and H98A/E62K variants were measured from wild-type and Lyn^-/-^ cells as described in (C), quantitated by densitometric analyses as fraction of VLPs secreted from wild-type set at 100%. Error bars represent mean±s.d. **(F)** VLPs (wild-type, H98A and H98A/E62K) were fractionated on sucrose and Optiprep gradients as described in Figure 6 to detect their subcellular distribution. **(G)** VLP-containing fractions co-migrating with Lyn on OptiPrep gradients described in (F) were scaled-up, concentrated, resolved by gel electrophoresis and detected by Coomassie staining for identification by mass spectrometry. **(H)** Entire lanes on gels were sliced into 1mm sections and subjected to trypsin digest. The peptide mix generated was processed and analysed by LTQ Orbitrap mass spectrometer. List of candidates were generated from peptide abundances isolated from VLP-containing fractions. **(I)** Control or VLP-secreting cells were stably transfected with GFP-RFP-LC3 reporter for measuring autophagic flux, and distinguish between formation of secretory versus degradative autophagosomes (*upper panel*). Control cells were cultured in complete media or starvation media to induce formation of degradative autophagosomes as controls (lower panel). **(J)** Formation of autophagosomes was measured by counting fluourescent puncta that were GFP+ and RFP+ (n = ∼1000 cells) and presented as abundance per cell. Autolysosomes were measured by quantitating RFP+ red puncta over ∼1000 cells. Error bars were calculated as mean±sd. **(K)** Schematic of secretion for immature versus processed mature virions through the conventional secretory pathway versus unconventional secretory organelles as determined by co-purifying proteins from mass spectrometry analyses.

Furin-sensitive VLPs secreted from wild-type cells displayed the anticipated pattern of cleaved pr and M fragments, whereas those carrying furin-resistant VLPs displayed a strong signal from uncleaved prM (**Figure 7B**). To determine whether any differences existed between transport kinetics of the VLP-variants, we collected them at different time intervals to quantitate the amount secreted (**Figure 7C**). Densitometric analyses revealed that secretion of furin-resistant immature VLPs was significantly slower compared to the processed, mature VLPs (**Figure 7D**). Given their distinct transport kinetics, we further assessed whether this was Lyn-dependent. Interestingly, whereas wild-type and furin-sensitive VLPs followed Lyn-dependent secretion, the furin-resistant immature VLPs displayed no secretion defect in Lyn^-/-^ cells, indicating that they followed a different transport mechanism (**Figure 7E**). To measure their subcellular distribution, we separated homogenates of VLP-producing cells on sucrose and OptiPrep gradients as described in Figure 6. Wild-type and furin-sensitive VLPs co-migrated with Rab11 and Tfr as seen with Dengue virions. However, furin-resistant VLPs were found enriched in the ER/ERGIC compartments instead (**Figure 7F**). To isolate the distinct VLP-containing vesicles, we concentrated the gradient fractions enriched in wild-type, H98A and H98A/E62K VLPs and resolved them by gel-electrophoresis (**Figure 7G**). The lanes were then sliced into 2 mm sections and subjected to trypsin digest followed by identification by mass spectrometry. Candidates identified in the wild-type and furin-sensitive VLP fractions represented factors previously implicated in the secretory autophagy/amphisome machinery ^29^, whereas those isolated with the furin-resistant VLPs reflected vesicles of the conventional secretory pathway (**Figure 7H**). To confirm that VLP-secretion was accompanied by formation of secretory autophagosomes, we generated control and VLP-secreting cells expressing a reporter for autophagic flux. RFP-GFP-LC3 cells appear as diffuse cytosolic fluorescence. Upon induction of autophagy, they appear as punctae - yellow, in autophagosomes and eventually red in autolysosomes upon degradation of the acid sensitive GFP signal. This reporter is therefore able to distinguish between generation of autophagosomes and its fusion with lysosomes for degradation. Using this reporter, we determined that control cells (not secreting VLPs) displayed basal levels of autophagosome and autolysosome formation (∼10-20/cell for each), whereas VLP-secreting cells contained a significantly higher proportion of autophagosomes (∼100/cell), which did not follow the lysosomal degradation route. In contrast when exposed to starvation conditions, these cells had an abundance of autolysosomes (>80/cell) (**Figure 7I, 7J**). These data support our mass spectrometry data that VLP secretion triggers formation of autophagosomes that are non-degradative. Collectively, these data indicate that mature VLPs trigger biogenesis of specialised secretory autophagosomes from the Golgi compartment. These organelles allow for efficient secretion of virions, and in the absence of Lyn, fuse with the lysosomal compartment for degradation. Immature virions on the other hand are unable to trigger this pathway and are instead exported through bulk exocytosis (**Figure 7K**). These results define an entirely new route for secretion and offer the possibility that infectious virions are secreted encapsulated in vesicles. These virions would therefore be equipped to evade circulating antibodies and furthermore dictate tissue tropism depending on their mode of entry.

## Discussion

In the current study we investigated the role of Src-family kinases (SFKs) in flavivirus secretion. SFKs were recently described to play a KDELR-dependent signalling role at the Golgi by triggering anterograde protein transport during increased flux through the secretory pathway ^30^. Since Dengue virus is known to hijack the KDELRs for trafficking to the Golgi, we hypothesised that a similar signaling cascade might be activated that facilitated virus secretion ^16^. Furthermore, SFK inhibitors, such as dasatinib and AZD0530 were previously reported to efficiently block Dengue infection by affecting virus assembly and secretion ^31^.

To test our hypothesis we screened for SFKs that were activated during virus infection. We identified three members – Src, Fyn and Lyn – that displayed increased phosphorylation in their activation loop in cells infected with Dengue or Zika. This phenomenon was recapitulated in cells that stably expressed the corresponding structural proteins prM and E, and constitutively secreted VLPs. Pharmacological inhibition of the SFKs significantly attenuated secretion, both in infected and VLP-producing cells. Among the identified SFKs, genetic depletion or deletion of Lyn caused the most dramatic defect. A combined depletion of Src and Lyn resulted in an additive effect.

We first confirmed that attenuated virus production in Lyn-deficient cells was specifically due to a block in secretion and not entry or replication. In cells deficient in Lyn, or expressing kinase-inactive or palmitoylation-deficient mutants, entry and replication remained unaffected, as did kinetics of transport from the ER to the Golgi compartments. On the other hand post-Golgi secretion was disrupted, resulting in a significant loss of E-protein arrival at the plasma membrane, and appearance of progeny virions/VLPs in the supernatants. This defect could be rescued when Lyn^-/-^ cells were reconstituted with wild-type Lyn but not an enzymatically inactive or palmitoylation-deficient mutant.

To further analyse the specific block in virus transport sustained in Lyn-deficient cells, we biochemically separated organelles of the secretory/endolysosomal pathway from infected cells to measure how Lyn and virus particles partitioned. In wild-type cells, both Lyn and virions were enriched in membranes that were positive for Rab11 and the transferrin receptor. These markers have been reported to be enriched in both recycling endosomes as well as exosomal and secretory autophagosomal vesicles ^27, 29, 32^.

Contrary to wild-type Lyn, upon expression of the kinase or palmitoylation mutants, virions were enriched in the lysosomal fractions. Inhibiting lysosomal degradation could rescue a significant fraction of viral E-protein. Interestingly, this mode of Lyn-dependent transport was specifically triggered by processed, mature virions. Exocytosis of protease-resistant immature virions occurred with significantly slower kinetics, and in a lyn-independent manner, suggesting that infectious virus particles might trigger a separate route for exiting cells, as compared to bulk secretion.

Current evidence on mechansims of flavivirus secretion are sparse, especially at the post-Golgi steps. Class-II ADP-ribosylation factors and KDELRs have been described for Dengue transport at a pre-Golgi step ^13, 33^. Although several components of the endosomal sorting complex (ESCRTs) were identified in Dengue and Japanese Encephalitis virus infection, they were found to participate in budding of immature virus particles into the ER-lumen rather than at the later stages of secretion ^34^. More interesting observations have emerged from Zika infection, where EM studies have identified large vesicles containing multiple virions, as well as small vesicles containing individual virions *en route* to fusion at the plasma membrane ^35, 36^. Our data from VLP-enriched fractions suggest that progeny virions exploit secretory autophagosomes to exit host cells and require Lyn activity to do so. Several outstanding questions remain, including the identity of Lyn substrates that are phosphorylated and recruited for virus transport, which will be addressed in future studies.

S-palmitoylation is a reversible post-translational modification that can regulate protein function in a manner similar to that of ubiquitylation and phosphorylation. Palmitoylation of soluble proteins often regulates their membrane association, intracellular trafficking and domain localisation. The process of de-/repalmitoylation constitutes a spatially organizing system that generates a directional flow from the Golgi to the PM, and counteracts the tendency to increase entropy by protein redistribution over all membranes ^23^. Palmitoylation of Lyn is thus necessary to prevent aberrant membrane targeting and lysosomal degradation of the subset of virus containing vesicles.

We have limited knowledge on how flaviviruses restructure the secretory pathway to generate membrane-bound compartments for viron assembly and release. Although the basic secretory machinery is conserved in all eukaryotic cells, it is likely that virion transport and release may proceed differently depending on cell types, with contributions from multiple parallel pathways. Flaviviruses infect a wide range of cells. Notable amongst them are monocytes, which have a highly expanded trafficking pathway for exosomal vesicles. Whether they function to transport flaviviruses in infected monocytes and the involvement of Lyn in the process, merits further investigation that will be pursued in future studies.

## Materials and Methods

### Cells Lines, Viruses and Antibodies

The following cell lines were used in this study: BHK21, Vero E6, HeLa, and HepG2 cells obtained from ATCC. Cells were maintained in DMEM or EMEM supplemented with 10% fetal bovine serum (FBS) and 1% penicillin/streptomycin at 37°C, with 5% CO2. The stable cell lines expressing prME-DENV1-4, prME-ZIKV or Lyn WT/C468A/C3S mutants were established using the retroviral vector pCHMWS-IRES-Hygromycin, selected following a 2-week period in the presence of 500 µg/mL hygromycin and maintained thereafter in the same medium, as previously described^37^. For biochemical analyses, the following antibodies were used: mouse anti-E mAb 4G2 was prepared using hybridoma cells D1-4G2-4-15 from ATCC; rabbit anti-phosphorylated Src family mAb, rabbit anti-Lyn mAb, rabbit anti-Src mAb and rabbit anti-Fyn from Cell Signaling Technology; mouse anti-GAPDH mAb from Abcam. Phospho-Tyrosine Mouse mAb (P-Tyr-100) conjugated magnetic bead from Cell Signaling Technology.

Virus stocks were prepared for DENV2 (16681), DENV1 (Hawaii), DENV2 (New Guinea), DENV3 (H87), DENV4 (Jamaique 8343), ZIKV (MR766) by determining tissue culture infective dose 50% (TCID_50_/ml) in Vero E6 cells challenged with 10-fold serial dilutions of infectious supernatants for 90 min at 37°C. Cells were subsequently incubated in DMEM with 2.5% FCS.

### Virus infections

#### RT-qPCR Assay to measure virus infection

Cells were plated in 96-well plates and infected at an MOI of 5. Infected cells were collected at indicated time intervals (as specified in the figure legends). For quantitation of RNA, cells were washed first with PBS and collected in 250µl of Trizol reagent for isolation of total RNA. Real time PCRs were performed with one-step or two-step methods using SYBR green or Taqman chemistry with gene specific primers.

#### Plaque assays

Serial dilutions of supernatants from infected cells were performed by adding on to BHK21/Vero E6 monolayers. After adsorption for 60 min at 37°C, cells were washed and plaque media was overlaid on the cells. After 3-6 days of incubation at 37°C, the monolayers were stained with crystal violet and plaques were counted.

### Screen for phosphorylated SFKs by DENV infection

3 × 10^6^ Vero E6 or BHK21 cells were seeded a day before infection. Attached cells were challenged by DENV1 with an MOI of 5 and harvested at 1day post infection. Harvested cells were lysed on ice with 1ml RIPA buffer (1% Triton X-100, 150 mM NaCl, 50 mM Tris-HCl, (pH 7.5), 1 mM EDTA, 0.5% Na-deoxycholate) supplemented with freshly added protease inhibitors cocktail (Roche) and phosphatase inhibitor tablets (Roche) for 30 min. Immunoprecipitation was performed on conjugated anti-phospho-SFKs antibodies and eluates were separated by gel electrophoresis and visualised by sliver staining. Entire lanes were sliced into 2-mm sections, and were further processed in 50% water/methanol. Samples were trypsinized and subjected to an LTQ Orbitrap mass spectrometer for identification of candidates. MS/MS spectra were analyzed using Sequest algorithm searching a composite target-decoy protein sequence database. The target sequences comprised the human protein repository of the Uniprot database. Decoy sequences were obtained upon reversing the orientation of target sequences. Allowed criteria for searches required trypsin cleavage (two missed cleavages allowed), peptide mass tolerance of 20 p.p.m, variable oxidation of methionine residues, and static carbamylation modification of cysteine residues. Peptide-spectrum matches were determined with estimated false discovery rate<1%. Spectral counts for each condition were combined at a protein level and normalized by protein length to infer protein abundances and intensities in each case. Identified hits were further categorized into different biological pathways using Gene Ontology analyses and Ingenuity Pathway Analyses software.

### Immunoprecipitation (IP) assay

Cell lysates (CL) were pre-cleared by incubation with 30µl of 50% Protein G Sepharose beads (Amersham Pharmacia Biotech) for 1 hour. Pre-cleared lysates from mock or DENV infected cells were then incubated for overnight at 4°C with 30µl of 50% Protein G Sepharose beads conjugated with either p-SFKs (for phosphorylated SFKs screen) or P-Tyr-100 conjugated magnetic beads (to validate selected SFK candidates). Subsequently, beads were collected by centrifugation at 13,000 rpm for 30 sec at 4°C and washed three times with cold wash buffer (50mM Tris-HCl buffer, pH 7.4, containing 0.1% (wt/vol) Triton X-100, 300mM NaCl, 5mM EDTA) supplemented with 0.02% (wt/vol) sodium azide and phosphatase inhibitor tablets, and once with cold PBS. Bound proteins were eluted by boiling in 30 µl 2×SDS-PAGE loading buffer, separated by gel electrophoresis, and visible in silver stain or analysed by Western blotting using appropriate antibodies.

### Luminex assay

MILLIPLEX MAP 8-Plex Human SFK Phosphoprotein Kit was purchased from Millipore and used according to the manufacturer’s instructions. Gapdh MAPmate kit was used as loading control. Cell lysates from virus-infected cells were collected and protein concentrations adjusted to 1mg/ml. 25µl of each lysate preparation was mixed with 8-Plex SFK bead set and Gapdh beads per well of a 96-well plate. Following incubation overnight at 4°C on a slow shaker, 25µl of 1X biotinylated antibody mixture was added to each well. After incubating for 1h at room temperature with slow shaking, 25 µl of 1X streptavidin-phycoerythrin solution was added to each well. After incubating for 15 min at room temperature, 25µl of amplification buffer was added to each well and incubated for 15 min at room temperature. The reaction mix was resuspended in 150µl of assay buffer and analysed on Luminex 200^TM^ analyser. Median fluorescence intensities (MFI) for triplicate wells were averaged and normalised to average MFI of Gapdh as loading control.

### siRNA experiments

All siRNAs used in this work, including non-targeting (NT) siRNA (D-001206) and transfection reagents DharmaFECT1 (T-2001) were purchased from Dharmacon. Src siRNA (L-003175), Fyn siRNA (L-003140) and Lyn siRNA (L-003153) were provided as SMARTpool ON-TARGET plus siRNAs, which are pools of four siRNAs targeting various sites in a single gene. For siRNA experiments, reverse transfection was performed using DharmaFECT1 reagents as recommended by the manufacturer. Briefly, siRNAs mixed with DharmaFECT1 reagents were added to 24-well plates in DMEM medium without FBS and antibiotics. Twenty minutes later 0.8 ml cells (60,000 cells/ml in DMEM supplemented with 10% FBS) were added to each well to final siRNA concentrations. Cells were then incubated at 37°C for 72 hours. For VLP assays, medium was replaced with 0.3 ml of Opti-MEM and, 14 hours later, culture supernatant (SN) containing secreted VLPs was collected and clarified by centrifugation at 4,000 rpm for 5 min. Cells separated from supernatants were lysed in RIPA buffer containing freshly added protease inhibitors cocktail (Roche), for 30 min on ice. For DENV infection assay, siRNA treatment was performed 48 hours before viral infection using the same conditions described above, and the siRNA-transfected cells were re-seeded to avoid over-confluency before virus challenge. Cells and supernatants containing progeny viruses were collected and processed as described above.

### VLP quantification

To detect VLP secretion, 90µl of supernatants and cell lysates from prME expressing stable cell lines or transiently transfected cells were added with 30µl 4X NuPAGE LDS sample buffer and subjected to western blotting using anti-E antibody 4G2. The mean luminescence and area of E protein signals detected were measured by densitometry using Image Quant TL (Thermo Fisher Scientific Inc) and analysed by image J software.

### SU6656 treatment

Src family Kinase inhibitor-SU6656 was purchased from Selleckchem and dissolved in DMSO to make a 50mM stock. Cell cytoxicity was first measured by MTT assay. To test VLP secretion in SU6656 treated cells, 2 × 10^5^ DENV1-4 prME expressing cells were pre-seeding a day before treatment. SU6656 with indicated concentrations as specified in figures was added to cells. Control cells (0 µM) received the same amount of DMSO. After 6 h treatment, medium was changed into 500µl Opti-MEM supplemented with SU6656 or DMSO. Supernatants and cell lysates were collected 16hrs later to measure secreted versus total VLPs respectively.

### CRISPR/Cas9 mediated deletion of Lyn

Potential target sequences for CRISPR interference were found using the rules outlined elsewhere ^38^. SgRNA targeting of Lyn was designed and cloned into the chimeric CRISPR/Cas9 vector PX459. Upon confirming potential off-target effects of the seed sequence using NCBI human nucleotide blast, none of the bases overlapping with any other location of human genome was found. The PX459-sgRNA clone was used to transfect HeLa (for VLP assay), HepG2 and Huh7 cells (for infection assay). Cells were selected on puromycin and immunoblotted with anti-Lyn to verify deletion.

### Pulse-chase analyses of viral protein transport

Pulse-chase experiments were performed as previously described^39^. Briefly, ∼ 1 × 10^7^ cells (mock or virus-infected) were detached by trypsinization and starved for 30 min in cysteine/methionine free medium at 37°C prior to pulse labelling. Cells were labelled in 10 mCi/ml of [^35^S]Cys/Met for 10 min in a 37°C water bath, and chased in cold medium for indicated time intervals. At each time point, aliquots were withdrawn and the reaction stopped with cold PBS. Cell pellets were either stored for further processing or labelled with NHS-biotin for surface labelling. For measuring transport characteristics of E-protein, cell pellets were lysed in Tris buffer containing 0.5% NP-40. Lysates were pre-cleared with agarose beads for 1 hr at 4°C, followed by immunoprecipitations for 3 hr at 4°C with end-over-end rotation. Immunoprecipitated samples were eluted by boiling in reducing sample buffer and subjected to SDS-PAGE followed by detection using autoradiography.

### Lck inducible expression

Lentiviral particles were produced in HEK293T cells by co-transfection of Lck lentiviral expression vectors with the packaging plasmids as described elsewhere ^40^. Viral supernatants were collected after 48 h, filtered and used for transduction of cells with 5 µg/ml Polybrene. Puromycin selection was applied at 1 µg/ml for Lck expression.

### Statistical analysis

Results are presented as mean±s.d of the specified number of experiments. Comparisons between two populations of data were made using the Student’s unpaired *t*-test with a confidence limit for significance set at 0.05 or less.

### Biochemical fractionations for membrane separation of the secretory pathway

Mock and virus-infected cells were pelleted and washed three times with PBS. Cells were homogenised in 10 mM Tris buffer (pH 7.2) containing 400mM sucrose, 1 mM EDTA supplemented with protease inhibitor cocktail, using a loose pestle Dounce homogeniser. Homogenates were centrifuged at 100,000 × g for 45 min to collect total membranes or centrifuged sequentially at 1000 × g (10 min), 3000 × g (10 min), 25,000 × g (20 min) and 100,000 × g (30 min; Beckman TLA100.3 rotor) to collect membrane fractions. The 25,000 × g and 100,000 × g fractions were combined and resuspended in 750 µl of 1.25 M sucrose buffer and overlaid with 500 µl of 1.1 M and 500 µl of 0.25 M sucrose buffer. Centrifugation was performed at 120,000 × g for 2 hr (Beckman TLS 55 rotor), after which the interface between 0.25 M and 1.1 M sucrose was collected and resuspended in 1 ml 19% OptiPrep for a step gradient containing 0.5 ml 22.5%, 1 ml 19% (sample), 0.9 ml 16%, 0.9 ml 12%, 1 ml 8%, 0.5 ml 5% and 0.2 ml 0% OptiPrep each. The gradient was centrifuged at 150,000 × g for 3 hr (Beckman SW 55 Ti rotor) and subsequently ten fractions, 0.5 ml each, were collected from the top.

## Acknowledgements

The authors acknowledge Nicla Porciello for help with Lck plasmids. This work was supported by Health and Medical Research Funds (16150592 and 16150732), the Scientific Research Plan of the Beijing Municipal Education Committee (KM201710025002), and BNP Paribas CIB. SS is supported by the Croucher Foundation.

