## Supplementary information for "Activation of Src-family kinases orchestrate secretion of flaviviruses by targeting mature progeny virions to secretory autophagosomes"

**Expression of acylated Lck is able to rescue Lyn-deficiency during Dengue infection**

Within the SFK members, Lyn, Lck and Hck comprise the SrcB subfamily, displaying significant overlap in homology, fatty acylation and subcellular distribution (S. J. Parsons and J. T. Parsons, 2004b; Senis et al., n.d.). To determine whether a related member within this subfamily can rescue Lyn-deficiency, we engineered Vero cells to stably express inducible Lck in the $Lyn^{-/-}$ background. Cells expressing the wild-type, palmitoylation-deficient mutant (C3S) and an N-terminal truncation (ΔSH4) that cannot be myristoylated or palmitoylated were generated by doxycycline-inducible lentiviral transduction. All the Lck constructs could be expressed in the $Lyn^{-/-}$ cells in an inducible manner as verified by immunoblotting (**Figure S4A**). We challenged these cells with Dengue virus and measured viral titers. Wild-type Lck but not the acylation-deficient variants (C3S or ΔSH4) could rescue viral titers in the $Lyn^{-/-}$ cells, as quantitated by plaque assays (**Figure S4B**). In parallel, secretion alone was measured in cells expressing the corresponding VLPs. Again, wild-type Lck but not the acylation-deficient variants (C3S or ΔSH4) could rescue the secretion defect in $Lyn^{-/-}$ cells quantitated by the appearance of E-protein in supernatants (**Figure S4C**). To confirm the effect of Lck on the subcellular distribution of virus particles, we performed biochemical fractionations of secretory organelles from homogenates of Dengue-infected cells (**Figure S4D**). Reminiscent of Lyn activity, wild-type Lck and viral E-protein co-migrated with markers of recycling endosomes and plasma membrane. On the other hand, in the presence of the C3S or ΔSH4 mutant, Lck and viral E-protein co-sedimented with lysosomal fractions. These data indicate that SFK members with similar biophysical properties and membrane affinity can facilitate virus secretion. However, only a partial rescue of virus production could be achieved with Lck in Zika-infected cells, suggesting that specific interactions between viral proteins and the SFKs might be necessary in transport of progeny virions (**Figure S5**).

**Materials and Methods:**

**MTT assay to determine cell viability**

$2\times10^4$ parental HeLa/Vero E6 and the corresponding DENV1-4 prME expressing cells were pre-seeding in 96 well plates. After overnight incubation, 50mM SU6656 stock was diluted in cell culture medium to a final concentration as indicated in figures and added to cells. For 0μM control, same volume of DMSO as that in the SU6656 treated group was added to cells. After a day of incubation, cell culture medium was replaced with 200μl 0.8mg/ml MTT solution (Sigma) followed by incubation for 4hrs. Finally, the formazan crystals formed by the MTT reagent was dissolved in 100 μl of isopropanol (Merck) and incubated at 4ºC for 30 minutes of shaking. The percentage of viable cells was calculated by comparing the absorbance of SU6656 treated groups to the 0μM control group at 570 nm.

**2-Bromopalmitate treatment to inhibit palmitoylation**

Each 60-mm plate of VLP-secreting cells was starved for 1 h in DMEM containing 2.5% FBS and 0.5% defatted BSA with or without indicated concentrations of 2-bromopalmitate. After replacing with fresh media and overnight culture at 37 °C, the VLPs were collected from supernatants, concentrated by ultracentrifugation, subjected to SDS-PAGE, and immunoblotted with anti-E 4G2 antibodies.

**siRNA depletions of SFKs**

Src siRNA (L-003175), Fyn siRNA (L-003140) and Lyn siRNA (L-003153) were provided as SMARTpool ON-TARGET plus siRNAs, which are pools of four siRNAs targeting various sites in a single gene. For siRNA experiments, reverse transfection was performed using DharmaFECT1 reagents as recommended by the manufacturer. Briefly, siRNAs mixed with DharmaFECT1 reagents were added to 24-

well plates in DMEM medium without FBS and antibiotics. Twenty minutes later 0.8 ml cells (60,000 cells/ml in DMEM supplemented with 10% FBS) were added to each well to final siRNA concentrations. Cells were then incubated at 37°C for 72 hours. For VLP assays, medium was replaced with 0.3 ml of Opti-MEM and, 14 hours later, culture supernatant (SN) containing secreted VLPs was collected and clarified by centrifugation at 4,000 rpm for 5 min. Cells separated from supernatants were lysed in RIPA buffer containing freshly added protease inhibitors cocktail (Roche), for 30 min on ice.

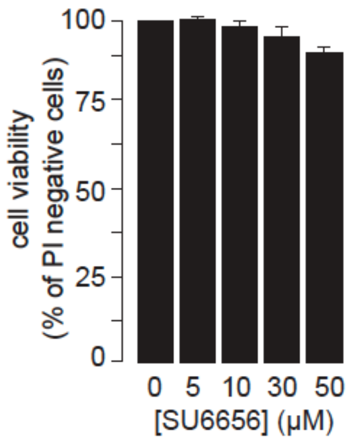


**Figure S1: Cell viability upon treatment with SU6656 inhibitor**

Cells were treated with varying concentrations of the selective SFK inhibitor SU6656 and incubated overnight at 37ºC, following which, culture medium was replaced with 200μl 0.8mg/ml MTT solution and incubated for 4hrs. Formazan crystals formed by the MTT reagent were dissolved in 100 μl of isopropanol and incubated at 4ºC for 30 minutes of shaking. The percentage of viable cells was calculated by comparing the absorbance of SU6656 treated groups to the 0μM control group (DMSO only) at 570 nm.

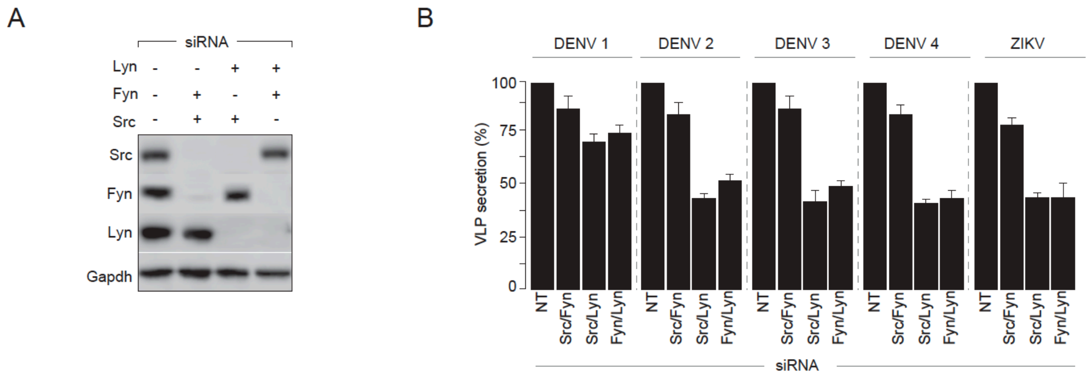


**Figure S2: Combined gene-depletions of SFKs inhibit VLP secretion**

**(A)** Combinations of Lyn, Fyn and Src depletions were performed by co-transfections of siRNA targeting individual kinases in VLP-producing cells, and verified by immunoblotting with specific antibodies. **(B)** VLP secretion from non-targeting (NT) and siRNA-treated cells was measured using 4G2 antibodies to detect viral structural proteins as readout. Quantitation of VLP secretion was performed by densitometric analyses for all serotypes of Dengue and Zika virus. Percentage secreted was calculated as fraction of total intracellular VLPs normalised to untreated cells set at 100%. Error bars represent mean±s.d from atleast three independent biological replicates.

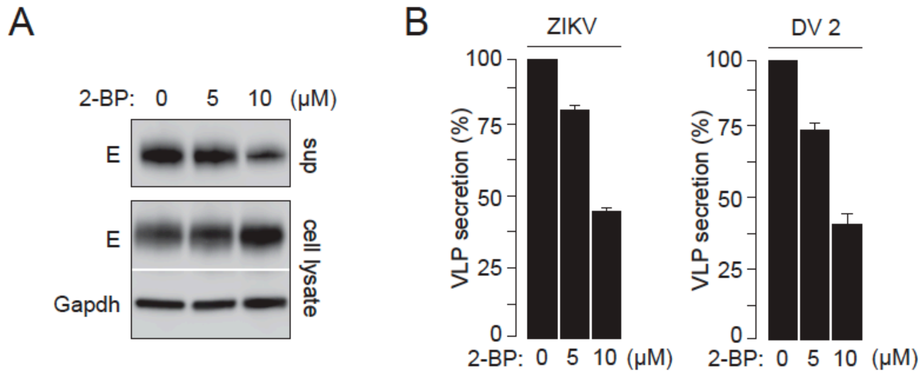


**Figure S3: Inhibition of palmitoylation with 2-bromopalmitate impairs VLP secretion**

**(A)** VLP-producing cells were treated with 0, 5 or 10 μM 2-BP and cultured overnight at 37ºC. Secreted VLPs were collected from supernatants, concentrated, resolved by SDS-PAGE and detected by anti-E 4G2 antibodies. Corresponding lysates were prepared to measure total intracellular VLPs by gel electrophoresis and detection by immunoblotting using 4G2 antibodies. Gapdh levels were measured as loading controls. **(B)** Quantitation of VLP secretion was performed by densitometric analyses for Dengue 2 and Zika virus. Percentage VLP secreted was calculated as fraction of total intracellular VLPs normalised to untreated cells set at 100%. Error bars represent mean±s.d from atleast three independent biological replicates.

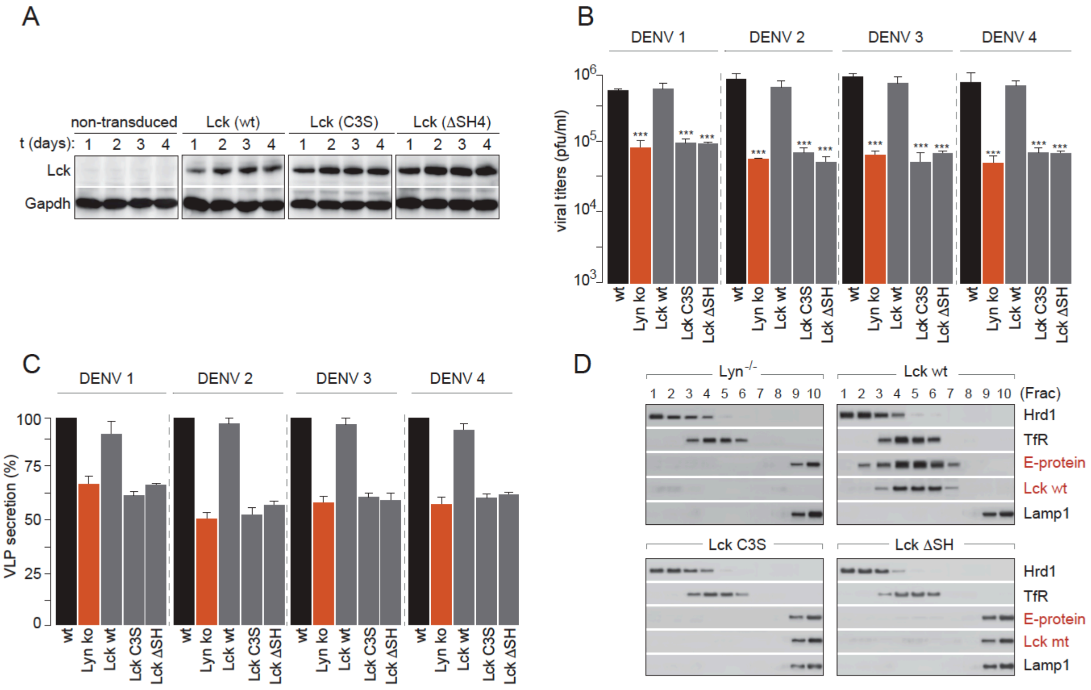


**Figure S4. Lyn-dependent Dengue virus transport from the Golgi can be rescued by expressing Lck**

**(A)** Inducible expression of wild-type, palmitoylation-deficient (C3S) and SH4-domain truncation (ΔSH4) mutants of Lck in the Lyn$^{-/-}$ cells. Validation of Lck expression was performed upto 4 days post induction by immunoblotting with Lck-specific antibodies. **(B)** Lck-expressing cells described in (A) were challenged with Dengue at MOI 2. Supernatants collected from infected cells were tested for viral titers using plaque assays. Error bars represent mean±s.d from three independent biological replicates; Statistical significance was calculated using Student's T test; *** represents p<0.001. **(C)** VLP-secretion was measured in Lck-expressing cells by immunoblotting for viral structural proteins in supernatant preparations. **(D)** Subcellular distribution of viral E-protein in membrane fractions were generated from Lck-expressing cells by biochemical separations described in figure 6B.

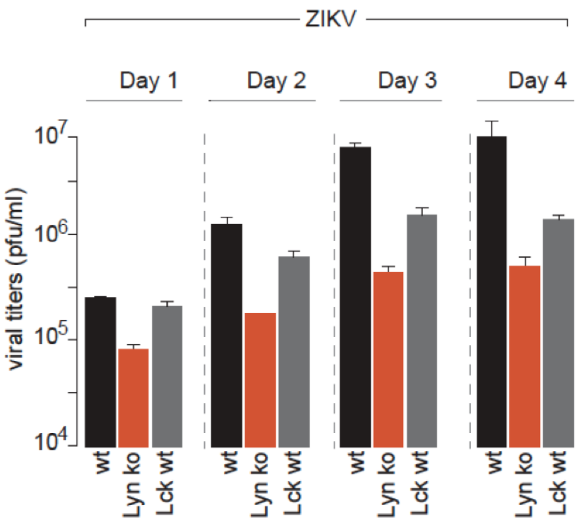


**Figure S5: Partial rescue of Zika virus production by Lck-expression in Lyn$^{-/-}$ cells**

Inducible expression of wild-type Lck in the Lyn$^{-/-}$ cells were performed as described in Figure 7. Lck-expressing cells were challenged with Zika at MOI 2. Supernatants collected from infected cells at indicated time points were tested for viral titers using plaque assays. Error bars represent mean±s.d from three independent biological replicates.
